# A site-resolved post-translational modification framework for drug-response analysis

**DOI:** 10.64898/2026.11.19.767872

**Authors:** Moustafa Zein, Aboul Ella Hassanien

## Abstract

Post-translational modifications (PTMs) regulate protein activity and treatment-associated signalling, providing functional information that can connect drug-response estimates with candidate pharmacodynamic measurements. Assessing this information requires a framework that determines its contribution beyond fixed molecular features while retaining the identity of the modified sites underlying model predictions. We developed the PTM Biological Dynamics Layer (PTM–BDL), which integrates typed PTM states with protein sequence, structure-derived and drug chemical features to predict log-transformed half-maximal inhibitory concentration and resistance while providing site-level interpretation through integrated gradients. Retrospective evaluation used source-derived and propagated PTM profiles across biologically distinct BCR–ABL, HDAC/EP300 and EGFR/ERBB2 settings, yielding internal areas under the receiver operating characteristic curve (AUROCs) between 0.785 and 0.946. Retraining demonstrated a consistent incremental contribution from PTM inputs in K562/CML, with a mean AUROC increase of 0.0268 and positive gains in all five folds, alongside more modest incremental gains in HeLa/HDAC and EGFR/ERBB2. Within K562/CML, CRKL Y207 and STAT5A Y694 retained the leading attribution ranks for their proteins across three seeds, while greater PTM attribution in dasatinib than in cytotoxic drug contexts identified a focused setting for pharmacodynamic investigation. These findings support context-dependent use of PTM information and connect predictive assessment with prioritisation of individual sites, providing a basis for targeted pharmacodynamic measurements and independent experimental validation.

## 1. Introduction

Drug response depends on whether treatment alters the cellular processes that sustain survival and proliferation. The presence of a drug target does not by itself specify this dependence. Downstream signalling can persist despite inhibition of the intended target, as illustrated by resistance to ERBB-directed therapies (Arteaga & Engelman, 2014). Protein sequence, structure, and compound chemistry provide molecular context but do not fully specify the regulatory state of a protein in a particular cell or after treatment. Post-translational modifications (PTMs) are relevant to this distinction because they regulate protein activity, interactions, localisation, and stability (Narita et al., 2019; Seet et al., 2006). Relating PTM states to drug response therefore offers a way to examine functional information about targets and their signalling networks alongside their molecular identities.

The systems considered here illustrate complementary biological reasons for including PTMs. In BCR–ABL signalling, CRKL phosphorylation provides a readout of kinase activity and its inhibition by imatinib (Hamilton et al., 2006; Hantschel, 2012). In an HDAC/EP300 setting, phosphorylation and acetylation are relevant to chromatin regulation (Fischle et al., 2003), while non-histone acetylation regulates protein activity, stability, and interactions (Narita et al., 2019). For ERBB receptors, N-glycosylation can affect receptor stability and activation (Fujitani et al., 2024). Interfering with this process has restored inhibitor sensitivity in experimental models of EGFR-inhibitor resistance (Lopez Sambrooks et al., 2018). These findings motivate examining PTMs both as regulators of drug-relevant biology and as readouts of pathway modulation. Site identity is also important because different modified residues can participate in different regulatory mechanisms (Ochoa et al., 2020).

Baseline PTM states and treatment-associated changes address distinct pharmacological questions. Baseline profiles can characterise the signalling state in which a drug acts. Changes after exposure can report how that state responds to treatment. Evidence for both uses already exists. DRUML uses omics-derived response-distance features to rank anticancer drugs (Gerdes et al., 2021), and proteomic and phosphoproteomic measurements have supported ex vivo response modelling in acute myeloid leukaemia (Gosline et al., 2022). MPhos used diagnostic phosphoproteomic profiles to predict midostaurin-plus-chemotherapy response in independent cohorts (Borek et al., 2024). Acute phosphoproteomic changes after drug exposure have also predicted sensitivity and combination effects in non-small-cell lung cancer (Coker et al., 2022). These studies establish the biological and predictive relevance of PTM-related information, motivating evaluation of its contribution in the specific drug response setting under study.

The question addressed here is when site-specific PTM states contribute response-relevant information beyond fixed protein and compound features, and which sites account for that contribution. Predictive accuracy alone does not answer this question because a model can exploit molecular context without making substantial use of the PTM inputs. Addressing it requires retaining the biological distinctions between modification types, sites, baseline states, and treatment-associated changes, then evaluating their contribution to the response endpoint. This also differs from predicting whether a residue can be modified or how a compound changes PTM abundance. The DrugPTM-Bench, for example, concerns drug-induced, cell-type-specific PTM changes (Badkul et al., 2026). Here, harmonised PTM profiles are inputs to drug response analysis. The research need is to connect response estimates with an assessment of the value of these regulatory-state inputs and with specific sites that can be examined in pharmacodynamic assays.

We developed the PTM Biological Dynamics Layer (PTM–BDL) to address this question. A separate PTM branch preserves protein, site, and modification-type identity together with baseline and change values. Type-conditioned gating and masked self-attention accommodate different PTM classes and variable site panels before fusion with fixed sequence, structure-derived, and chemical features. This separation enables comparison of models trained with and without PTM information, while integrated gradients for individual sites identify the inputs on which a fitted prediction depends (Sundararajan et al., 2017). Attention provides a candidate way to represent dependence among sites. Classical models can also use features for individual sites and provide attribution, so the architecture requires evaluation against those alternatives. Inputs in this study combine source-derived and propagated profiles. Propagation does not constitute a new measurement of the recipient cell–drug context.

Recent benchmarks reinforce the need to distinguish the biological question from architectural complexity. DrEval found that complex drug response models did not outperform properly tuned tree ensembles in relevant evaluation settings (Bernett et al., 2026). In genetic-perturbation transcriptome prediction, metric calibration revealed deep-learning performance that conventional aggregate metrics could obscure (Miller et al., 2026). These findings motivate strong baselines and evaluation aligned with the intended scientific use. We evaluate response prediction, incremental PTM contribution, and attribution behaviour separately.

Our working hypothesis is that PTM states can provide complementary information about drug-relevant protein regulation, with a contribution that varies across biological settings. We examine this hypothesis retro-spectively in BCR–ABL-associated phosphorylation, HDAC/EP300-associated phosphorylation and acetylation, and EGFR/ERBB2-associated phosphorylation and glycosylation. The objective is to determine when the available PTM profiles contribute predictive information beyond fixed molecular features and which individual sites merit pharmacodynamic investigation. The framework combines comparisons against matched classical baselines with retraining, controlled input removal, and assessment of attribution stability. These complementary analyses separate the added predictive contribution of PTM profiles from the influence of individual inputs on fitted predictions.

## 2. Materials and methods

### 2.1. Study design and data sources

We conducted a retrospective, multimodal modelling study in three biological settings represented by K562/CML, HeLa/HDAC, and EGFR/ERBB2. These case-study names identify the reference signalling systems and PTM source contexts. The response cohorts include additional cell lines and are not experiments confined to K562 or HeLa. The intended user is a researcher examining how configured PTM inputs affect a response model and selecting sites for subsequent matched assays. This is a method-characterisation study using existing response data, not a prospective screening or clinical validation study. The unit of analysis was a cell-line–drug–target-protein context, with a continuous natural-log-transformed half-maximal inhibitory concentration (lnIC50) and a binary resistance label. IC50 values are expressed in micromolar units before the natural logarithm is taken. The GDSC processing pipeline labels ln(IC_50_) ≥ 0, corresponding to IC_50_ ≥ 1 *μ*M, as resistant and values below this cutoff as sensitive. Curated entries retain their recorded labels. This is an operational in vitro classification, not a clinically established resistance threshold for each drug. Drug response data from public datasets and published studies were harmonised for model training and evaluation. Endpoints were obtained primarily from the Genomics of Drug Sensitivity in Cancer resource (Yang et al., 2013). The EGFR/ERBB2 cohort also used DepMap mutation annotations (Tsherniak et al., 2017) to assign receptor context. Canonical protein sequences and site nomenclature were taken from UniProt. Representative experimental structures were selected from the RCSB Protein Data Bank (Burley et al., 2019). Literature-derived response values supplemented cell–drug pairs absent from the selected GDSC records. PTM evidence was assembled from DrugPTM-Bench, case-specific perturbational phosphoproteomic studies, and, for the EGFR/ERBB2 case, glycoproteomic and EGFR-resistance studies supporting site selection and interpretation (Badkul et al., 2026; Fujitani et al., 2024; Lopez Sambrooks et al., 2018; Tozuka et al., 2024).

The cohort accounting is reported in Supplementary Table S1. In the HeLa/HDAC and K562/CML cases, each endpoint was expanded over two and three target-protein contexts, respectively. These rows therefore share an outcome label and must not be treated as independent drug response experiments. The EGFR/ERBB2 cohort contained 943 GDSC endpoints and eight additional response endpoints. The HeLa/HDAC cohort contained 1,676 GDSC and 11 additional endpoints, and the K562/CML cohort contained 3,544 GDSC and eight additional endpoints.

### 2.2. Problem formulation

For a context consisting of a protein sequence *s*, representative structure *q*, drug structure *m*, and a configured set of *n* PTM sites, the model takes 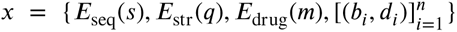. Here, *b_i_* is the harmonised baseline value for site *i* and *d_i_* is its harmonised treatment-associated change. The two outputs are *ŷ*, an lnIC50 prediction, and *p̂*, the probability of the recorded resistance class. This formulation distinguishes a baseline-only input configuration from one that additionally contains a treatment-associated PTM change. The latter addresses retrospective pharmacodynamic associations. The baseline-only analysis reported here removes the change channel from an already fitted model. It is not a separately validated prospective predictor. Treatment-associated values would require an assay or another explicitly validated source before use in a new context.

### 2.3. Data harmonisation and PTM-site encoding

Each case study used a fixed, biologically curated site schema. The EGFR/ERBB2 table contains phosphorylation and glycosylation slots. The HeLa/HDAC table contains phosphorylation and acetylation slots, and the K562/CML table contains phosphorylation slots. Raw baseline intensities, when present on a large intensity scale, were transformed as log_2_(1 + *x*). Small-scale values were retained. Source-derived per-site drug profiles were calculated as mean treatment-associated log2 fold changes after peptide-to-protein site mapping.

The resulting values are not uniformly direct measurements for the particular cell line, drug, protein, and site represented by a row. Where an exact match was unavailable, source profiles were propagated. In the EGFR/ERBB2 case, profiles were transferred within a mutation or ERBB2 amplification class, with an attenuation factor of 0.85 for propagated fold changes and additional cell-line modifiers based on mutation context. In the HeLa/HDAC and K562/CML cases, source baselines were adjusted by tissue-specific multipliers and drug profiles by drug-specific factors (0.1–1.0). Thus, these features are more accurately described as *evidence-informed, harmonised PTM states* than as uniformly measured per-row PTM observations.

Missing configured baseline values are represented by the reference level *b_i_* = 1 and missing treatment-associated values by *d_i_* = 0. The same reference state is used in the no-PTM counterfactual. This is a model input convention, not a biological assertion of wild-type occupancy. Each site also receives a configured PTM-subtype label. The registry distinguishes real sites from padding introduced to give proteins a common tensor width, and the PTM transformer excludes padded slots through its attention and pooling masks. Baseline-only inputs retain *b_i_* and set every *d_i_* to zero, whereas pharmacodynamic inputs retain both channels.

The pipeline assigns a propagation-confidence score to every row. It is a rule-based provenance score derived from source matching, tissue group, and drug class. It does not identify a direct per-row measurement, represent a calibrated probability, or enter the predictor as a learned feature. It is used only to define a high-provenance sensitivity analysis, in which PTM values with a score below 0.90 are reset to a reference state. The score documents the strength of the PTM match, but it cannot validate a propagated value. Supplementary Table S2 reports the resulting provenance and PTM-vector-diversity accounting for each case study.

### 2.4. Multimodal feature extraction

#### 2.4.1. Protein-sequence representation using ESM-2

Protein representations were extracted before supervised training with the pretrained ESM-2 650M protein language model (Z. Lin et al., 2023). Per-residue embeddings were retained and adaptively pooled to at most 200 tokens where necessary to control the cost of joint self-attention. The supervised predictor reads these arrays from disk. ESM-2 is not an encoder module within the trained predictor and is not fine-tuned.

#### 2.4.2. Fixed structure-derived graph features

The structural arrays are also fixed during predictor training, but they must not be described as pretrained structural embeddings. For EGFR/ERBB2, a 512-dimensional, three-layer graph encoder is constructed from PDB residue identity, B-factor, spatial distance, and sequence separation, without pretrained weights. The HeLa/HDAC and K562/CML arrays use the same non-pretrained graph-feature path. Hence, all three modalities are fixed inputs to PTM–BDL, whereas only the sequence and chemical arrays are outputs of pretrained encoders.

#### 2.4.3. Drug representation using ChemBERTa

ChemBERTa token-level and pooled chemical arrays (Ahmad et al., 2022) were extracted from canonical drug structures before supervised training. They are read as fixed 384-dimensional inputs. As for ESM-2, the ChemBERTa weights are not part of the optimised predictor.

### 2.5. PTM–BDL architecture

Figure 1 presents the prediction workflow and site-token construction. The implementation settings are specified below. Attribution procedures are described in Section 2.8.

**Figure 1:**
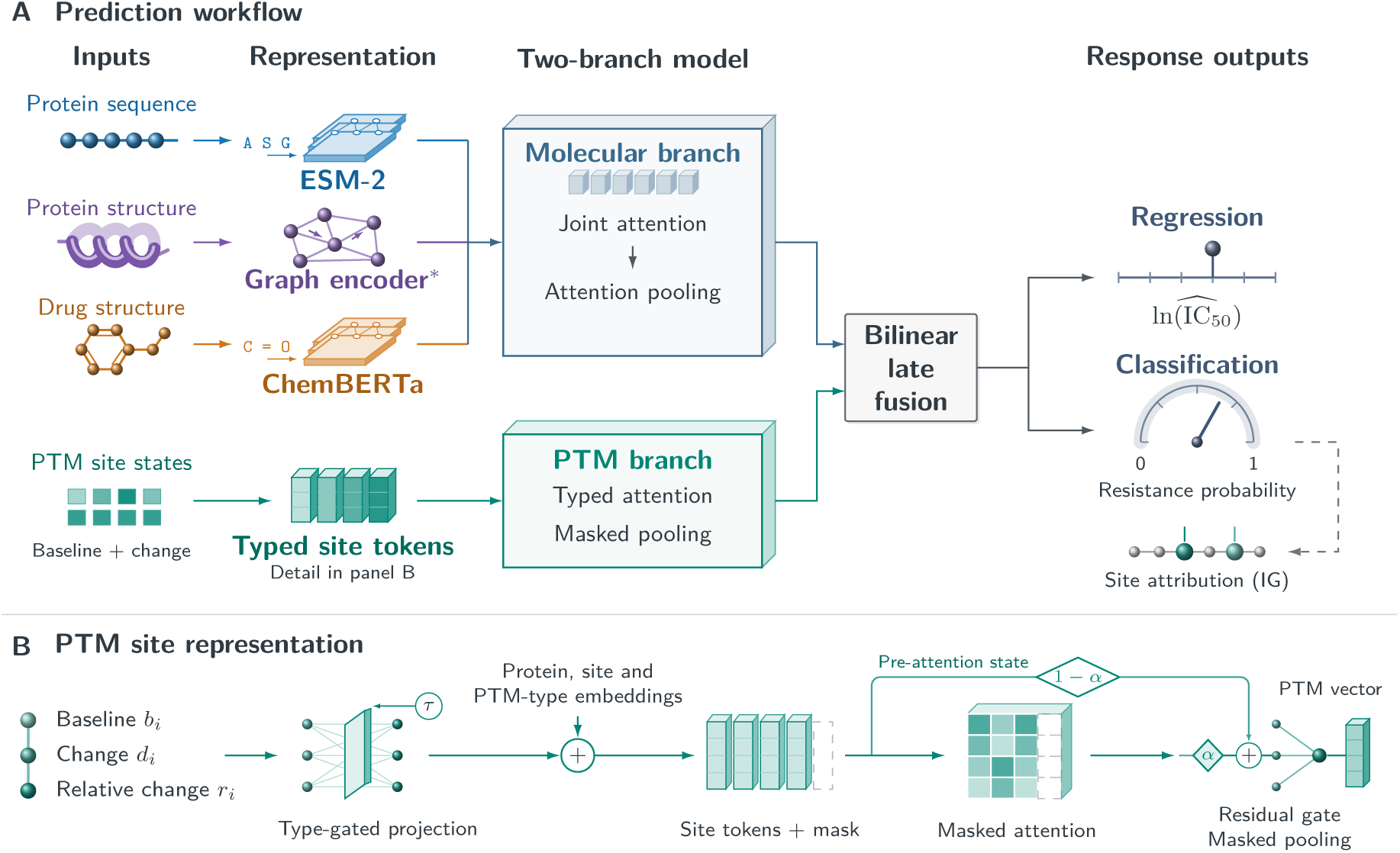
Post-translational modification biological dynamics layer (PTM–BDL). (A) Blue, purple, orange, and teal identify sequence, structure, drug, and PTM inputs, respectively. ESM-2 and ChemBERTa provide fixed features from pretrained protein and chemical language models; the asterisk marks the non-pretrained structure encoder. Molecular and typed-site representations are combined by bilinear late fusion to estimate lnIC50 (the natural logarithm of the half-maximal inhibitory concentration expressed in *μ*M) and resistance probability. (B) Site *i* supplies baseline *b_i_*, treatment-associated change *d_i_*, and relative change *r_i_* = *d_i_*∕( *b_i_* + 10^−6^). Projection is conditioned on PTM type *τ*; protein, site-slot, and PTM-type embeddings are then added. After masked attention, the learned gate *α* weights the attended state and 1 − *α* weights the pre-attention state carried by the bypass. Their sum undergoes masked mean pooling to form the PTM vector. Circled plus signs denote vector addition; unfilled tokens and attention-grid cells indicate excluded padding. Solid arrows show computation; the dashed arrow leads to post-hoc integrated gradients (IG) for site attribution. Inputs include source-derived and propagated PTM profiles.

#### 2.5.1. Static molecular branch

The static branch independently projects the 1,280-dimensional sequence, 512-dimensional structure-derived, and 384-dimensional chemical token embeddings to 512 dimensions. Modality embeddings are added, the three token streams are concatenated, and a four-layer, eight-head transformer produces a joint representation. Gated attention pooling then yields a 512-dimensional static-context vector.

#### 2.5.2. PTM biological dynamics branch

The numerical triplet in Fig. 1B is projected through two linear layers (3 to 64 to 64 dimensions), with layer normalisation, Gaussian error linear unit (GELU) activation, and dropout between them. A sigmoid-activated linear gate takes the concatenated projected values and PTM-subtype embedding. Learned subtype, target-protein, and site-slot embeddings each have 64 dimensions, preserving site and modification identity alongside the numerical state.

#### 2.5.3. Typed self-attention

The PTM transformer has two four-head layers followed by layer normalisation. Residual-gate coefficients are computed by a sigmoid-activated linear layer from the concatenated attended and pre-attention states. The registry supplies the real-site mask for attention and pooling. The final PTM representation has 64 dimensions. Attention can represent dependencies between PTM types when more than one type is configured, but its weights are descriptive model quantities and are not interpreted as causal effects.

#### 2.5.4. Bilinear late fusion

Separate linear projections of the static and PTM representations are passed through tanh, multiplied element-wise, and transformed by a 512-dimensional fusion multilayer perceptron.

#### 2.5.5. Dual prediction heads

Two task-specific multilayer heads share the fused representation and are trained jointly with the objective in Section 2.6. The classification head returns a logit, converted to probability by a sigmoid for inference. The outputs are related summaries of sensitivity, rather than independent biological endpoints.

### 2.6. Training objective and optimisation

For the recorded internal experiments, training used class-balanced sampling. Validation and test sets were not resampled. The EGFR/ERBB2 configuration used a learning rate of 10^−4^ and weight decay of 10^−5^. HeLa/HDAC and K562/CML used 10^−3^ and 10^−2^, respectively. The batch size was 16, dropout was 0.1, the maximum was 100 epochs, and early stopping used a patience of 15 epochs.

Models were optimised with AdamW and cosine learning-rate annealing. The multitask objective was *L* = *L*_Huber_ + 2*L*_focal_, with focal-loss parameters *α* = 0.25 and *γ* = 2 (T.-Y. Lin et al., 2017). Classification label smoothing was set to 0.05. Gradients were clipped at an *L*_2_ norm of 1.0. The checkpoint with the largest of validation AUROC and balanced accuracy was retained. A resistance threshold was subsequently selected only on the validation set by maximising Youden’s *J* and carried forward to test-set classification summaries.

### 2.7. Experimental design

#### 2.7.1. Data splitting

Rows were split 70%/15%/15% into training, validation, and test sets with seed 42, stratified by the joint target-protein and resistance-label category. This is an interpolation split, not a test of unseen cell–drug generalisability. In particular, the internal splits retain 326 shared cell–drug pairs between train and test in HeLa/HDAC and 1,298 in K562/CML because each endpoint was expanded across protein contexts. They also retain repeated PTM vectors across partitions. We therefore describe random-split performance as an internal contextual estimate only.

#### 2.7.2. Machine-learning baselines

Classical baselines comprise random forest, XGBoost, ridge, and elastic-net models. They use the same rows and splits as PTM–BDL, with mean-pooled sequence, structure-derived, and drug embeddings concatenated with the PTM features. Hyperparameters are selected by inner five-fold cross-validation on the training partition. Each classifier selects its own Youden-*J* threshold on the validation partition. These are matched tabular baselines, not independent reimplementations of published deep drug response systems. Published-number comparisons are retained only as contextual literature comparisons and are not treated as head-to-head experiments.

#### 2.7.3. Evaluation metrics

AUROC, area under the precision–recall curve for the sensitive class, balanced accuracy, sensitivity, specificity, and macro-F1 quantify the binary task. Root mean squared error and Pearson correlation quantify lnIC50 regression. Brier score, expected calibration error (ECE), and reliability diagrams assess calibration of resistance probabilities. ECE uses ten probability bins.

#### 2.7.4. Statistical testing

We calculated 95% confidence intervals using 1,000 bootstrap resamples. We also performed paired AUROC comparisons, Wilcoxon signed-rank tests for paired summaries, and Benjamini–Hochberg adjustment across recorded families of *p* values. The paired AUROC comparison used bootstrap resampling rather than the analytic DeLong covariance. It is therefore reported as a bootstrap-based paired AUROC test. Resampling is at the context-row level, not clustered by cell–drug endpoint. The five-fold Full-versus-No-PTM comparison uses a paired *t* test on fold-wise AUROC values. These folds share training observations, and endpoint expansion introduces dependence between contexts. The reported intervals and *p* values are therefore descriptive of the implemented resampling, not independent confirmation of biological PTM effects.

#### 2.7.5. Ablation configurations

The primary ablation protocol holds the fitted model, test split, and all non-ablated inputs fixed, then replaces the selected input with its defined reference state. PTM levels are set to one and treatment-associated changes to zero. Drug and structure arrays are zeroed. Configured PTM-type ranges are reset, and baseline-only, delta-only, provenance-restricted, and randomised-input conditions are evaluated. This controlled input-removal design tests the sensitivity of the fitted predictor to the specified replacement, without introducing changes in optimisation or random initialisation. The related concept of model reliance quantifies dependence on inputs (Fisher et al., 2019). Our reference-state replacement is distinct from its permutation-based estimator.

The evaluated configurations are No PTM, No Drug, No Structure, individual PTM-type removal or retention (glycosylation or acetylation where configured), baseline-only, delta-only, high-provenance PTM-only, randomised PTM, and typed self-attention versus an MLP PTM encoder. The no-typed-attention comparison is retrained across seeds 42, 123, and 456 to test the value of interaction processing, rather than the value of PTM state itself. Separately, a five-fold Full-versus-No-PTM experiment trains the two models within every fold. The retrained comparison uses stratified context-row folds and tests whether a PTM-associated difference persists after re-optimisation. It is reported separately from input replacement because the latter can also introduce an input distribution shift. Neither protocol establishes causal site effects.

#### 2.7.6. Generalisation protocols

Fresh models are trained for leave-one-drug-out, five-fold cold-cell, and leave-one-group-out evaluation. Within each training partition, an inner validation split selects the checkpoint and classification threshold. The held-out cells, drugs, or groups are used only once for final evaluation. GDSC-to-CTRP transfer is conducted separately for each case study. A strong generalisation claim requires all rows sharing a cell–drug endpoint to remain in the same partition and to be supported by the cold-cell and cold-drug results. These protocols are needed to distinguish learned response associations from memorisation of repeated endpoints or propagated PTM templates.

### 2.8. Interpretability analysis

#### 2.8.1. Integrated Gradients

We used integrated gradients to assign the prediction to PTM baseline and delta inputs (Sundararajan et al., 2017). The reference state is *b_i_* = 1, *d_i_* = 0 for every valid slot, matching the reference state used by the no-PTM input replacement. Attributions were computed along a linear path from this state to the observed harmonised state, aggregated by site and PTM channel, and summarised within protein and drug strata. They identify inputs to which the fitted model is sensitive. They do not demonstrate that a site causally mediates drug response. Attention maps are treated likewise as descriptive internal signals, not mechanistic proof.

#### 2.8.2. Cross-type attention analysis

For multi-PTM cases, we extract post-softmax attention from the final PTM transformer layer and average it over evaluated contexts after applying the real-site mask. We summarise within-type and between-type blocks, such as phosphorylation–acetylation and phosphorylation–glycosylation. These values describe internal model interactions and are interpreted alongside integrated gradients for individual sites to nominate cross-PTM interaction hypotheses.

#### 2.8.3. Joint PTM-class removal

For each two-type case study, we calculated a fixed-model inclusion–exclusion contrast on held-out AUROC. Let *A*(⋅) denote AUROC under the indicated reference-state replacement. The non-additive joint-reliance residual was

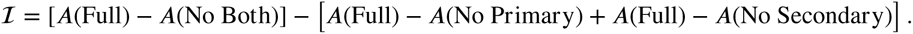

All removal arms use the same production model, test contexts, and PTM reference state (level = 1, delta = 0). A positive 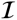 indicates that the joint removal reduces fitted-model discrimination more than the sum of the individual removals. We interpret this as non-additive cross-PTM model reliance, complementary to cross-type attention. It is a computational interaction signature to be tested experimentally.

#### 2.8.4. Attribution-stability analysis

For seed-specific models, we recompute integrated gradients for individual sites and compare rankings with pairwise Spearman correlation, top-3 and top-5 Jaccard overlap, and agreement of the top-ranked site. The stability analyses use three seeds (42, 123, and 456). Stability characterises reproducibility of the model explanation, not causal validity of a highlighted site.

### 2.9. Reproducibility and implementation

The implementation uses Python 3.11, PyTorch 2.1 or later, and scikit-learn. The analyses were run on a MacBook Pro with an Apple M1 Pro chip (10 CPU cores) and 16 GB unified memory using Apple MPS. ESM-2 650M and ChemBERTa-77M-MTR are the frozen checkpoints used to create the feature arrays. The trained predictor contains approximately 15.8 million trainable parameters, depending on the case-specific PTM registry. Split seed 42 is used for the internal experiments, and seeds 42, 123, and 456 for the multi-seed architecture and attribution analyses. The five-fold Full-versus-No-PTM retraining runs took 70.6, 219.5, and 835.6 minutes for EGFR/ERBB2, HeLa/HDAC, and K562/CML, respectively, at 30 epochs per fold. In the recorded EGFR/ERBB2 run, training took 2,100 s for PTM–BDL, 6.86 s for random forest, 69.81 s for XGBoost, 1.13 s for ridge, and 87.12 s for elastic net.

Case-specific site schemas, molecular identifiers, training settings, analysis scripts, dependency manifests, and a Docker configuration are provided in the source repository. Split indices, feature arrays, and checkpoints are generated as local analysis outputs. Data access is governed by the licences of GDSC, DepMap/CCLE, RCSB PDB, UniProt, and the PTM source studies.

## 3. Results

### 3.1. Internal response prediction and matched baselines

Internal predictive performance varied across the three biological settings (Table 1). PTM–BDL achieved AUROC values of 0.875 for EGFR/ERBB2, 0.946 for HeLa/HDAC, and 0.785 for K562/CML. The corresponding lnIC50 Pearson correlations were 0.597, 0.949, and 0.619, respectively. HeLa/HDAC had the highest AUROC and regression correlation, indicating that performance was sensitive to the biological setting and cohort composition.

**Table 1.** Internal context-split classification and regression performance of PTM–BDL and matched baseline models.

| Case study | Method | AUROC | AUPRC-sensitive | Balanced accuracy | Pearson *r* | RMSE |
|---|---|---|---|---|---|---|
| K562/CML | PTM–BDL | 0.785 | 0.801 | 0.751 | 0.619 | 2.320 |
|  | XGBoost | 0.814 | 0.826 | 0.750 | 0.652 | 2.195 |
|  | Ridge | 0.814 | 0.830 | 0.753 | 0.650 | 2.200 |
|  | Random forest | 0.810 | 0.823 | 0.751 | 0.648 | 2.206 |
|  | Elastic net | 0.812 | 0.828 | 0.752 | 0.645 | 2.212 |
| HeLa/HDAC | PTM–BDL | 0.946 | 0.956 | 0.935 | 0.949 | 1.077 |
|  | Random forest | 0.952 | 0.959 | 0.947 | 0.949 | 1.071 |
|  | XGBoost | 0.952 | 0.957 | 0.947 | 0.950 | 1.068 |
|  | Ridge | 0.942 | 0.955 | 0.938 | 0.950 | 1.061 |
|  | Elastic net | 0.942 | 0.953 | 0.932 | 0.950 | 1.063 |
| EGFR/ERBB2 | PTM–BDL | 0.875 | 0.667 | 0.822 | 0.597 | 1.800 |
|  | Elastic net | 0.927 | 0.730 | 0.794 | 0.720 | 1.391 |
|  | Ridge | 0.927 | 0.730 | 0.798 | 0.716 | 1.387 |
|  | XGBoost | 0.894 | 0.708 | 0.787 | 0.736 | 1.365 |
|  | Random forest | 0.888 | 0.662 | 0.710 | 0.650 | 1.560 |

Classical baselines achieved equal or higher AUROC on the same internal splits. The strongest baseline AUROCs were 0.927 for elastic net in EGFR/ERBB2, 0.952 for random forest in HeLa/HDAC, and 0.814 for ridge and XGBoost in K562/CML. The following analyses examine the incremental PTM contribution and the attribution and modality-reliance behaviour of the typed site-level representation.

The three systems also differed in calibration and cohort composition. The EGFR/ERBB2 classification scores had the largest ECE (0.442), whereas HeLa/HDAC and K562/CML had ECE values of 0.167 and 0.204, respectively. Per-drug, grouped, and cross-dataset analyses are consequently reported separately in Sections 3.4–3.6 rather than inferred from these aggregate scores.

### 3.2. Ablation and modality-contribution analysis

PTM inputs provided the largest improvement after retraining in K562/CML (Table 2). The mean AUROC difference between models trained with and without PTM inputs was 0.0057±0.0133 in EGFR/ERBB2 (*p* = 0.438), 0.0011 ± 0.0028 in HeLa/HDAC (*p* = 0.458), and 0.0268 ± 0.0132 in K562/CML (*p* = 0.015). The K562/CML difference was positive in all five folds. These findings indicate that the predictive contribution of the available PTM profiles varied across the biological settings. The nominal *p* values remain subject to the dependence described in Section 2.7.

**Table 2:**
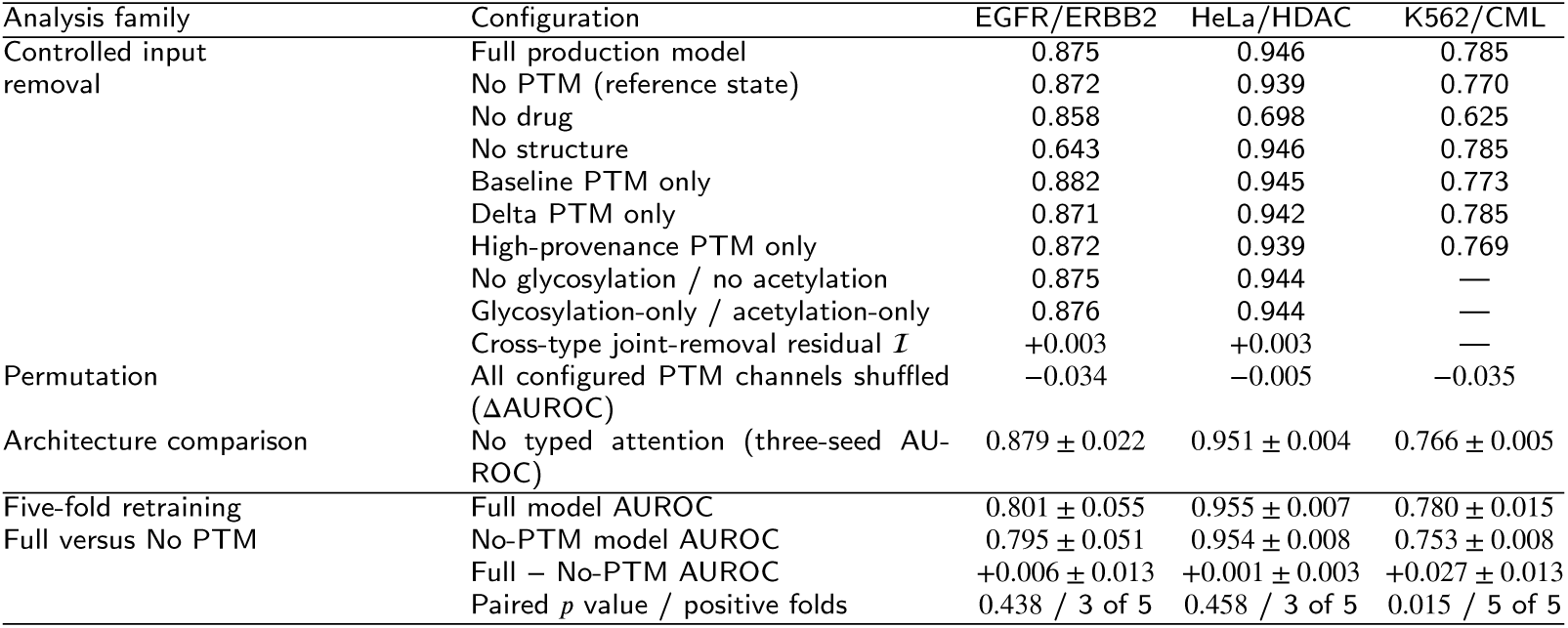
AUROC from controlled input removal, cross-type joint removal, and the retrained Full–No-PTM comparison. The No-PTM reference state sets PTM levels to one and treatment-associated changes to zero.

| Analysis family | Configuration | EGFR/ERBB2 | HeLa/HDAC | K562/CML |
|---|---|---|---|---|
| Controlled input removal | Full production model | 0.875 | 0.946 | 0.785 |
|  | No PTM (reference state) | 0.872 | 0.939 | 0.770 |
|  | No drug | 0.858 | 0.698 | 0.625 |
|  | No structure | 0.643 | 0.946 | 0.785 |
|  | Baseline PTM only | 0.882 | 0.945 | 0.773 |
|  | Delta PTM only | 0.871 | 0.942 | 0.785 |
|  | High-provenance PTM only | 0.872 | 0.939 | 0.769 |
|  | No glycosylation / no acetylation | 0.875 | 0.944 | — |
|  | Glycosylation-only / acetylation-only | 0.876 | 0.944 | — |
|  | Cross-type joint-removal residual ℐ | +0.003 | +0.003 | — |
| Permutation | All configured PTM channels shuffled (ΔAUROC) | −0.034 | −0.005 | −0.035 |
| Architecture comparison | No typed attention (three-seed AU-ROC) | 0.879 ± 0.022 | 0.951 ± 0.004 | 0.766 ± 0.005 |
| Five-fold retraining<br>Full versus No PTM | Full model AUROC | 0.801 ± 0.055 | 0.955 ± 0.007 | 0.780 ± 0.015 |
|  | No-PTM model AUROC | 0.795 ± 0.051 | 0.954 ± 0.008 | 0.753 ± 0.008 |
|  | Full − No-PTM AUROC | +0.006 ± 0.013 | +0.001 ± 0.003 | +0.027 ± 0.013 |
|  | Paired *p* value / positive folds | 0.438 / 3 of 5 | 0.458 / 3 of 5 | 0.015 / 5 of 5 |

The complementary input-removal analysis keeps the model and evaluation split fixed while replacing selected inputs with their defined reference states (Table 2). This distinguishes fitted-model reliance from the incremental predictive contribution assessed by retraining.

The controlled input-removal results reveal different modality profiles across the three case studies. Removing drug embeddings produced the largest AUROC reduction in HeLa/HDAC (0.248) and K562/CML (0.160), whereas removing the structure-derived input produced the largest reduction in EGFR/ERBB2 (0.232). Replacing all PTM values by the reference state changed AUROC by 0.003, 0.006, and 0.016 in EGFR/ERBB2, HeLa/HDAC, and K562/CML, respectively. Permuting all configured PTM channels reduced AUROC by 0.034, 0.005, and 0.035, indicating that the fitted models use the configured PTM arrangement to different degrees.

The baseline-only and change-containing configurations clarify which PTM state is used by each fitted predictor. Baseline-only AUROCs were 0.882, 0.945, and 0.773, whereas delta-only AUROCs were 0.871, 0.942, and 0.785 in EGFR/ERBB2, HeLa/HDAC, and K562/CML, respectively. Removing or retaining the secondary PTM channel produced little change in the EGFR/ERBB2 and HeLa/HDAC systems. The no-typed-attention MLP comparison was additionally retrained across three seeds. This is an architectural comparison of token-interaction processing. It does not test whether PTM state itself is informative. The separately retrained Full–No-PTM comparison reported above addresses that question.

The paired type-removal arms also provide a descriptive contrast for non-additive cross-PTM model reliance. The joint-removal residual 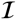 was +0.0029 AUROC for phosphorylation–glycosylation in EGFR/ERBB2 and +0.0028 for phosphorylation–acetylation in HeLa/HDAC. Thus, in both two-type systems the performance loss after resetting both classes exceeded the sum of the individual class-removal losses. These small residuals describe non-additivity of an AUROC-based input-replacement contrast. AUROC is itself nonlinear, so the contrast and cross-type attention do not isolate a molecular interaction or establish biological crosstalk. They are exploratory descriptors of the fitted model.

### 3.3. Site-level attribution and stability

Attribution analysis identified candidate PTM sites whose rankings varied in stability across the three settings. Integrated gradients quantified the absolute contribution of each configured site input to the resistance output on the recorded test contexts. Residue-level summaries exclude padding and unmapped auxiliary slots. EP300 phosphorylation outputs retain their recorded feature-column identifiers because these do not establish a consistent residue mapping.

In K562/CML, ABL1 Y264 had the highest mean absolute attribution among the recorded ABL1 phosphosites, whereas CRKL Y207 and STAT5A Y694 were the leading sites for their proteins in all three attribution seeds. CRKL phosphorylation provides a pharmacodynamic readout of BCR–ABL activity (Hamilton et al., 2006), and STAT5 activation has been implicated in BCR–ABL-driven leukemogenesis (Nieborowska-Skorska et al., 1999). These studies provide protein-level biological context for the rankings. ABL1’s leading site was less seed-stable, so its ranking is reported as a fitted-model observation, not as a claim that Y264 is a response-mediating site.

In the receptor system, EGFR Y1092 (Y1068 in mature-protein numbering) and ERBB2 Y1248 were the leading mean phosphosite attributions in the result artefact (Fig. 2a,b). These sites lie in receptor-signalling regions previously linked to adaptor-mediated signalling (Arteaga & Engelman, 2014; Schulze et al., 2005). The EGFR glycosylation channel was constant and had zero attribution, whereas ERBB2 glycosylation had non-zero mean attribution (Fig. 2c). Neither observation establishes a glycosylation mechanism. It records the information available to the fitted model.

**Figure 2:**
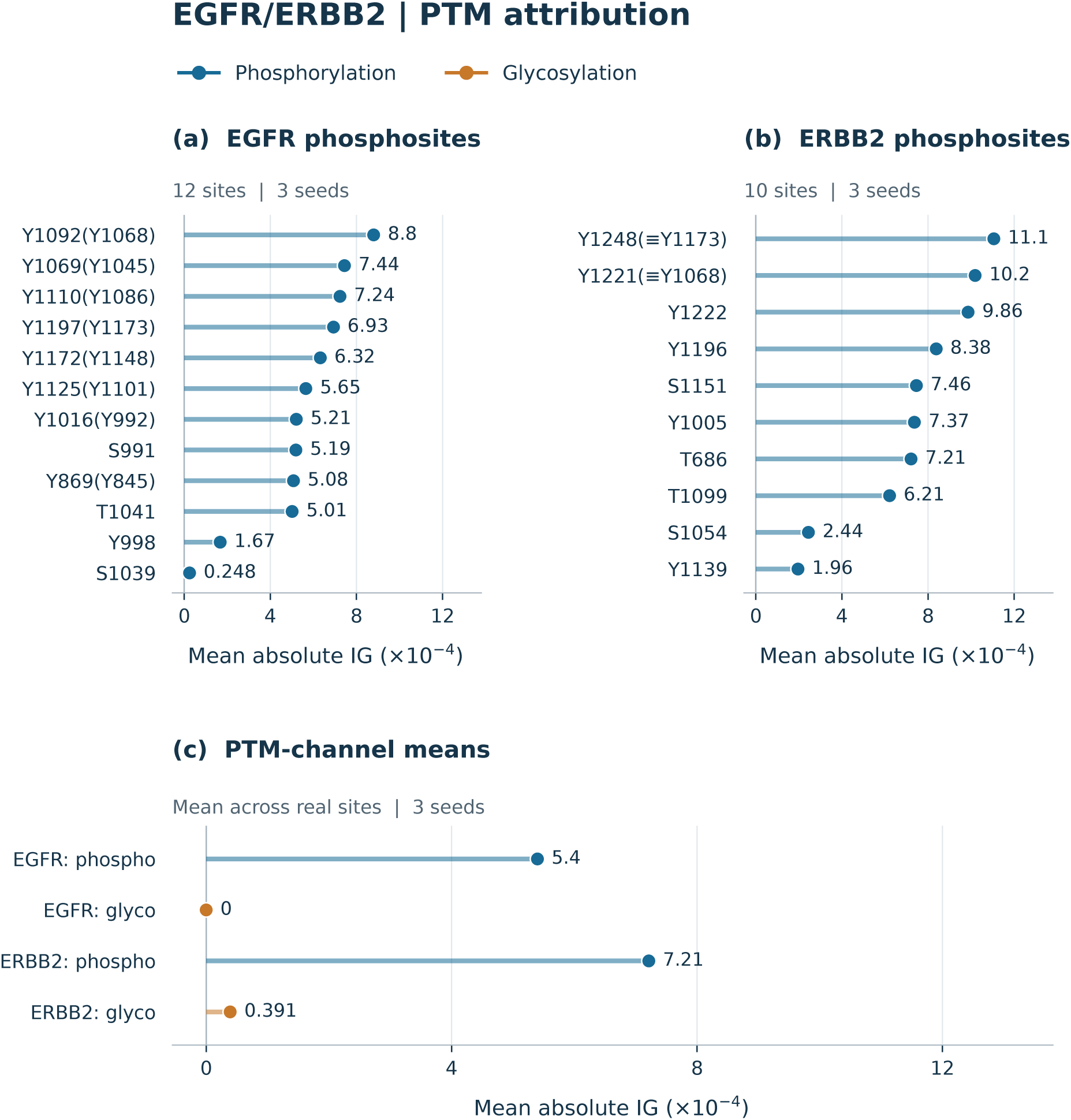
PTM attribution in the EGFR/ERBB2 case study. Panels (a) and (b) show mean absolute integrated gradients (IG), averaged across three seeds, for EGFR and ERBB2 phosphosites. Points are ordered by attribution magnitude; padding is excluded. Parenthetical labels indicate the recorded residue correspondences. Panel (c) shows the mean of site-level attributions within each protein–PTM channel. Blue denotes phosphorylation and orange denotes glycosylation. Numerical annotations use the axis multiplier, and zero values are retained.

In HeLa/HDAC, the result artefacts ranked the EP300 input labelled ptm_T445 and HDAC1 S393 as the leading phosphorylation features (Fig. 3a,b), and EP300 K1558 as the leading mapped acetylation feature (Fig. 3c). This demonstrates that the typed registry preserves phosphorylation and acetylation as distinct inputs. The EP300 phosphorylation identifier is a feature-column label, not evidence for a mapped EP300 T445 site. Their ordering varied between seeds, however, and HDAC1 acetylation could not be assigned to a site because the available channels were not site mapped. We therefore do not attach a biological interpretation to that channel.

**Figure 3:**
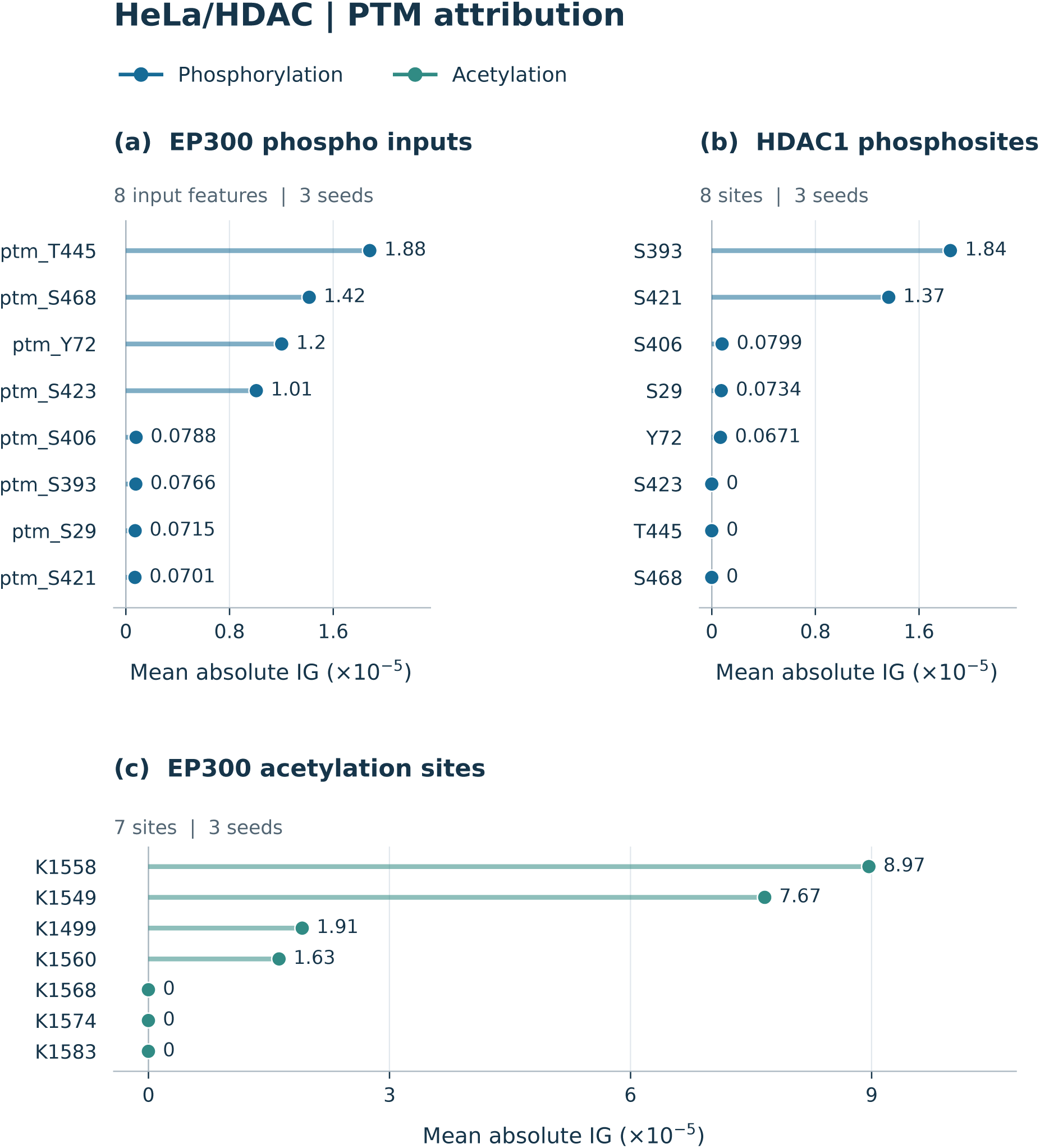
PTM-feature attribution in the HeLa/HDAC case study. Points show mean absolute integrated gradients (IG) across three seeds for EP300 phosphorylation inputs (a), HDAC1 phosphosites (b), and EP300 acetylation sites (c), ordered by magnitude within each panel. Blue denotes phosphorylation and teal denotes acetylation. EP300 phosphorylation labels identify input columns whose mapping to EP300 residues is unresolved. HDAC1 acetylation is omitted because its channels lack site mapping. Numerical annotations use the axis multiplier; panel (c) has a wider horizontal range than (a) and (b). Zero values are retained. Attribution magnitudes describe model sensitivity to the configured inputs.

### 3.4. Drug-dependent attribution patterns

The magnitude and distribution of PTM attribution differed between dasatinib and the cytotoxic drug contexts in K562/CML. Dasatinib (428 contexts) had a total PTM-attribution magnitude of 0.0877, with ABL1 Y89, Y185, and Y226 as its leading ABL1 sites. In contrast, the three chemotherapy groups had much smaller total magnitudes (0.00023–0.00045) and were led by ABL1 Y393 or Y412. Imatinib had only five test contexts and was therefore not used to draw a class-level conclusion.

These attribution differences identify kinase-inhibitor and cytotoxic-drug contexts for targeted comparison. Their magnitudes quantify sensitivity of the fitted prediction function, rather than the strength of a biological mechanism.

Cross-type analysis identifies a second, architecture-specific source of biological hypotheses. In HeLa/HDAC, the model assigned non-zero weights between phosphorylation and acetylation tokens, consistent with the known coupling of these regulatory layers (Fischle et al., 2003; Narita et al., 2019). EGFR and ERBB2 likewise showed reciprocal phosphorylation–glycosylation attention blocks, although the recorded sensitive–resistant differences were below 0.0005. These maps describe attention routing in the fitted model. Their small between-class differences and the weak secondary-channel removal effects limit their use for selecting a specific biological interaction.

### 3.5. Generalisation and transferability

Generalisation estimates were lower or more variable than the internal context-split estimates in several settings (Table 3). Cold-cell evaluation gave mean ± SD AUROCs of 0.815 ± 0.155 for EGFR/ERBB2, 0.957 ± 0.011 for HeLa/HDAC, and 0.771 ± 0.023 for K562/CML. By contrast, leave-one-drug-out AUROCs were 0.546 ± 0.220 across six evaluable EGFR/ERBB2 drugs, 0.428±0.053 across two evaluable HeLa/HDAC drugs, and 0.511±0.014 across five evaluable K562/CML drugs. The remaining HeLa/HDAC drug folds had a single test-set class, for which AUROC is undefined. The wide variation in the EGFR/ERBB2 cold-cell estimate and the lower drug-held-out estimates indicate that performance is conditional on the split and cannot be summarised by a single internal score.

**Table 3:** Cell-, drug-, and group-held-out performance, with the available external CTRP comparison. The number after each leave-one-drug-out result is the number of AUROC-evaluable held-out drugs. Checkpoint selection and threshold calibration for the rerun group-held-out analyses use only an inner validation split.

| Case study | Cold-cell AUROC | Leave-one-drug-out AUROC | Leave-one-group-out AUROC | CTRP comparison |
|---|---|---|---|---|
| EGFR/ERBB2 | 0.815 ± 0.155 | 0.546 ± 0.220 (6 drugs) | 0.548 ± 0.053 (2 groups) | *r* = 0.450 (*n* = 314) |
| HeLa/HDAC | 0.957 ± 0.011 | 0.428 ± 0.053 (2 drugs) | 0.899 ± 0.120 (3 groups) | *r* = 0.00002 (*n* = 986) |
| K562/CML | 0.771 ± 0.023 | 0.511 ± 0.014 (5 drugs) | 0.671 ± 0.131 (4 groups) | *r* = 0.002–0.120 (4 drugs) |

The leave-one-group-out analyses with inner-validation model selection produced mean AUROCs of 0.548 ± 0.053 in EGFR/ERBB2 (two evaluated groups), 0.899 ± 0.120 in HeLa/HDAC (three evaluated groups), and 0.671 ± 0.131 in K562/CML (four evaluated groups). Some groups were skipped because their held-out partition contained a single class. Reporting the number of evaluable groups makes that constraint explicit. Cross-group performance therefore varied across the evaluated case-study-specific groupings.

Agreement with external CTRP responses varied markedly across the three systems. EGFR/ERBB2 predictions had a Pearson correlation of 0.450 with CTRP outcomes across 314 matched contexts and four drugs. In HeLa/HDAC, the correlation was effectively zero (0.00002) across 986 Vorinostat contexts. For K562/CML, the model-to-CTRP correlations were 0.006 for Cytarabine (*n* = 615), 0.010 for Dasatinib (*n* = 1,542), 0.120 for Imatinib (*n* = 18), and 0.002 for methotrexate (*n* = 774). The Paclitaxel overlap contained 840 contexts but did not yield an estimable correlation. These results show heterogeneous transfer across the available settings. Table 3 summarises these evaluation protocols separately.

### 3.6. Robustness and calibration

Cross-validation retained the ranking of the three settings observed in the internal split. Mean ± SD AUROC was lower in EGFR/ERBB2 and K562/CML, at 0.821 ± 0.028 and 0.745 ± 0.010, respectively. HeLa/HDAC remained high at 0.953±0.008. The corresponding regression correlations were 0.495±0.038, 0.952±0.003, and 0.631±0.011 (Table 4). These estimates describe variability under the implemented context-row resampling scheme.

**Table 4:** Five-fold cross-validation classification and regression performance.

| Case study | AUROC | Balanced accuracy | Pearson *r* | RMSE |
|---|---|---|---|---|
| EGFR/ERBB2 | 0.821 ± 0.028 | 0.602 ± 0.058 | 0.495 ± 0.038 | 2.265 ± 0.141 |
| HeLa/HDAC | 0.953 ± 0.008 | 0.708 ± 0.092 | 0.952 ± 0.003 | 1.112 ± 0.077 |
| K562/CML | 0.745 ± 0.010 | 0.677 ± 0.057 | 0.631 ± 0.011 | 2.300 ± 0.058 |

Calibration was weakest in EGFR/ERBB2 (ECE 0.442, Brier score 0.253), compared with ECE/Brier values of 0.167/0.093 for HeLa/HDAC and 0.204/0.212 for K562/CML. The EGFR/ERBB2 bootstrap AUROC interval (0.767–0.987) was wide, which is consistent with its smaller test set. Reliability diagrams are provided separately for EGFR/ERBB2 (Supplementary Fig. S1), HeLa/HDAC (Supplementary Fig. S2), and K562/CML (Supplementary Fig. S3). Calibration and uncertainty estimates should therefore accompany, rather than be inferred from, AUROC. Attribution stability was heterogeneous. CRKL Y207 and STAT5A Y694 were the only recorded K562/CML top sites shared across all three seeds. Leading-site identities varied across seeds in EGFR/ERBB2 and HeLa/HDAC.

Excluding romidepsin from the HeLa/HDAC test set reduced AUROC from 0.946 to 0.505, while lnIC50 RMSE changed from 1.077 to 1.067. The fitted model and predictions were unchanged. Only the evaluated subset differed. This indicates that the high pooled classification performance depended strongly on the drug composition of the test set.

## 4. Discussion

### 4.1. Methodological contribution and pharmacological purpose

For pharmacological research, PTM–BDL provides a framework for assessing whether a selected PTM panel contributes useful information about drug response and for prioritising individual sites for subsequent measurement. The larger predictive contribution in K562/CML and the stable rankings of CRKL Y207 and STAT5A Y694 provide complementary evidence for prioritising matched pharmacodynamic assays in this setting. The small gains in HeLa/HDAC and EGFR/ERBB2 identify settings in which the available profiles contributed little additional predictive information. These findings connect model evaluation with specific questions about the utility of PTM measurements.

Prior work established the value of proteomic and phosphoproteomic response prediction (Borek et al., 2024; Coker et al., 2022; Gerdes et al., 2021). The present framework evaluates the incremental contribution of PTM inputs through retraining. Input removal and attribution provide complementary information about the sites and channels used by the fitted model. Preserving protein, site, and modification-type identity makes these findings traceable to candidates for pharmacodynamic follow-up.

### 4.2. Architectural rationale and comparison with simpler models

Matched classical baselines provide a strong reference for response prediction, consistent with DrEval’s finding that increased model complexity does not necessarily improve predictive performance (Bernett et al., 2026). The rationale for PTM–BDL is its structured representation of variable sets of typed sites. Gating, masked self-attention, and late fusion retain site information through a differentiable predictor that can be examined by integrated gradients. This design supports the analyses presented here, although simpler models can also support interpretation of individual sites.

The metric-calibration study of Miller et al. (2026) further motivates aligning evaluation with the biological signal of interest. That study examined transcriptomic responses to genetic perturbation. Its calibrated metrics and positive-control framework were not applied here. Brier score and ECE assess resistance-probability calibration, a separate concept. Support for PTM–BDL therefore rests on the observed context-dependent PTM contribution and its analysis of individual sites. The small gains in two settings and the additional computational cost favour selecting the architecture according to the analysis required, rather than assuming a general predictive advantage.

### 4.3. From site attribution to a pharmacodynamic hypothesis

CRKL Y207 and STAT5A Y694 were top-ranked for their proteins across three seeds. Published evidence for CRKL phosphorylation and STAT5 activation in BCR–ABL signalling supplies protein-level biological context (Hamilton et al., 2006; Nieborowska-Skorska et al., 1999). These rankings prioritise sites within a biologically selected panel. A focused follow-up is to assess whether measured changes at these sites track response across matched kinase-inhibitor concentrations and exposure times.

The larger total attribution for dasatinib than for the cytotoxic-drug contexts motivates testing whether the PTM panel is more informative in the kinase-inhibitor setting. Such assays would help distinguish measured biological variation from drug-specific propagation and model sensitivity. Variable rankings in the other systems, unmapped acetylation channels, and constant glycosylation inputs define where interpretation of individual sites is less resolved. Cross-type attention and residuals from joint removal similarly nominate model interactions for experimental examination.

### 4.4. Relationship to pharmacological methods research

Pharmacological methods studies connect a defined analytical problem with an appropriate validation setting. Examples include reliable negative sampling for drug–target prediction (Sharifabad et al., 2022) and interpretable toxicity modelling evaluated on external data (Masand et al., 2026). In the present study, the practical output is a response estimate accompanied by attribution to individual sites and input-replacement comparisons, helping an investigator organise candidate sites for targeted pharmacodynamic assays.

The current framework addresses retrospective response associations within the studied contexts. Pre-treatment application would require separate validation using baseline measurements. The present baseline-only analysis removes a channel from an already fitted model. Broader screening applications would also require stronger performance on unseen drugs and independent response datasets.

### 4.5. Limitations and future validation

Interpretation should account for how the PTM profiles represent the response contexts. Most rows use propagated profiles, with 0.1–2.3% meeting the rule-based high-provenance threshold, which is not equivalent to a direct matched measurement. PTM-associated gains may therefore reflect tissue, mutation, and drug context alongside site-state information. The small, biologically selected panels support focused analysis of the configured sites, with broader PTM coverage remaining to be evaluated.

Repeated endpoints and PTM profiles introduce dependence within the internal splits, row-wise cross-validation, and bootstrap resampling. The intervals and nominal *p* values should therefore be interpreted within these resampling schemes. Evaluations holding out cells, drugs, and groups address complementary transfer questions. The small number of evaluable drugs, limited unseen-drug discrimination, and variable external correlations constrain extrapolation beyond the studied settings. The operational resistance label and probability calibration, particularly in EGFR/ERBB2, also require assessment for any intended new use.

Input replacement can introduce distribution shifts, and stable attributions may reflect contextual associations as well as site-specific effects. Independent paired PTM and response measurements, with shared endpoints kept in the same partition, would strengthen assessment of added predictive value. Comparisons with interpretable classical models and matched time- and dose-resolved assays would help establish the value of the analyses of individual sites. The nominated sites and cross-PTM patterns remain hypotheses for such validation, rather than established causal mechanisms.

## 5. Conclusion

PTM–BDL connects drug response prediction with assessment of site-specific regulatory information and prioritisation of pharmacodynamic hypotheses. Across the retrospective case studies, retrained PTM-associated gains were largest in K562/CML and small in the other two systems, while classical baselines remained competitive or stronger. Stable CRKL Y207 and STAT5A Y694 rankings provide focused candidates for matched assays. The contribution is a framework that relates the predictive value of a PTM panel to candidates for pharmacodynamic measurement. Independent paired measurements and prospective evaluation are needed to establish broader predictive and biological value.

## CRediT authorship contribution statement

**Moustafa Zein:** Conceptualization, Methodology, Software, Data curation, Formal analysis, Investigation, Validation, Visualization, Writing – original draft, Writing – review and editing.

**Aboul Ella Hassanien:** Conceptualization, Formal analysis, Validation, Supervision, Writing – review and editing.

## Funding

This research did not receive any specific grant from funding agencies in the public, commercial, or not-for-profit sectors.

## Declaration of competing interest

The authors declare that they have no known competing financial interests or personal relationships that could have appeared to influence the work reported in this paper.

## Ethical statement

This study used publicly available data from GDSC2, CCLE/DepMap, UniProt, PDB, DrugPTM-Bench, and the cited source studies. No experiments involving human participants, human tissue, or animals were conducted. Ethical approval was therefore not required.

## Data availability

No new mass-spectrometry experiments or raw proteomics data were generated for this retrospective computational study. The primary data sources are publicly available from GDSC2, CCLE/DepMap, UniProt, RCSB PDB, and DrugPTM-Bench. The DrugPTM-Bench resource is derived from the decryptM ProteomeXchange dataset PXD037285. Source code, Docker support, and step-by-step instructions for generating processed features, splits, and model artefacts are publicly available at https://github.com/Moustafazn/PTM-BDL. Access to the underlying source data remains subject to the terms of the original data providers.

## Acknowledgements

We thank the GDSC, DepMap, UniProt, and PDB consortia for making their data publicly available. We are grateful to the DrugPTM-Bench team for providing the phosphoproteomic and acetylation benchmark data.

## Declaration of generative AI and AI-assisted technologies in the writing process

During the preparation of this work, the authors used ChatGPT (OpenAI) and Claude (Anthropic) to assist with language editing and improve code readability. The authors reviewed and approved the final manuscript.

## Supplementary material

**Table S1:** Case-study cohort accounting. Contexts are cell-line–drug–target-protein rows; HeLa/HDAC and K562/CML expand each cell–drug endpoint over two and three target proteins, respectively.

| Case study | Contexts | Cell–drug endpoints | Targets and drugs | PTM channels |
|---|---|---|---|---|
| EGFR/ERBB2 TKI | 951 | 951 | EGFR, ERBB2; six ERBB-family inhibitors | 12 phosphorylation and 12 glycosylation slots |
| HeLa/HDAC | 3,374 | 1,687 | HDAC1, EP300; six epigenetic modulators | 8 phosphorylation and 7 acetylation slots |
| K562/CML | 10,656 | 3,552 | ABL1, CRKL, STAT5A; two TKIs and three cytotoxic drugs | 12 phosphorylation slots |

**Table S2:** PTM provenance, vector diversity, and constant-channel accounting. The confidence threshold denotes high provenance under a rule-based source-matching heuristic; it does not establish a direct per-row measurement or estimate measurement uncertainty.

| **Data/provenance metric** | **EGFR/ERBB2** | **HeLa/HDAC** | **K562/CML** |
|---|---|---|---|
| *n* samples (train/val/test) | 665/143/143 | 2361/506/507 | 7458/1599/1599 |
| PTM unique vectors / *n* | 141 / 951 | 40 / 3,374 | 51 / 10,656 |
| PTM feature diversity | 0.148 | 0.012 | 0.005 |
| High-provenance (conf≥0.90) | 22 (2.3%) | 8 (0.2%) | 15 (0.1%) |
| Lower-provenance propagation | 929 (97.7%) | 3,366 (99.8%) | 10,641 (99.9%) |
| Constant PTM channels | 3 glyco Δ | 9 ph/ac Δ | 7 ph Δ |

**Figure S1:**
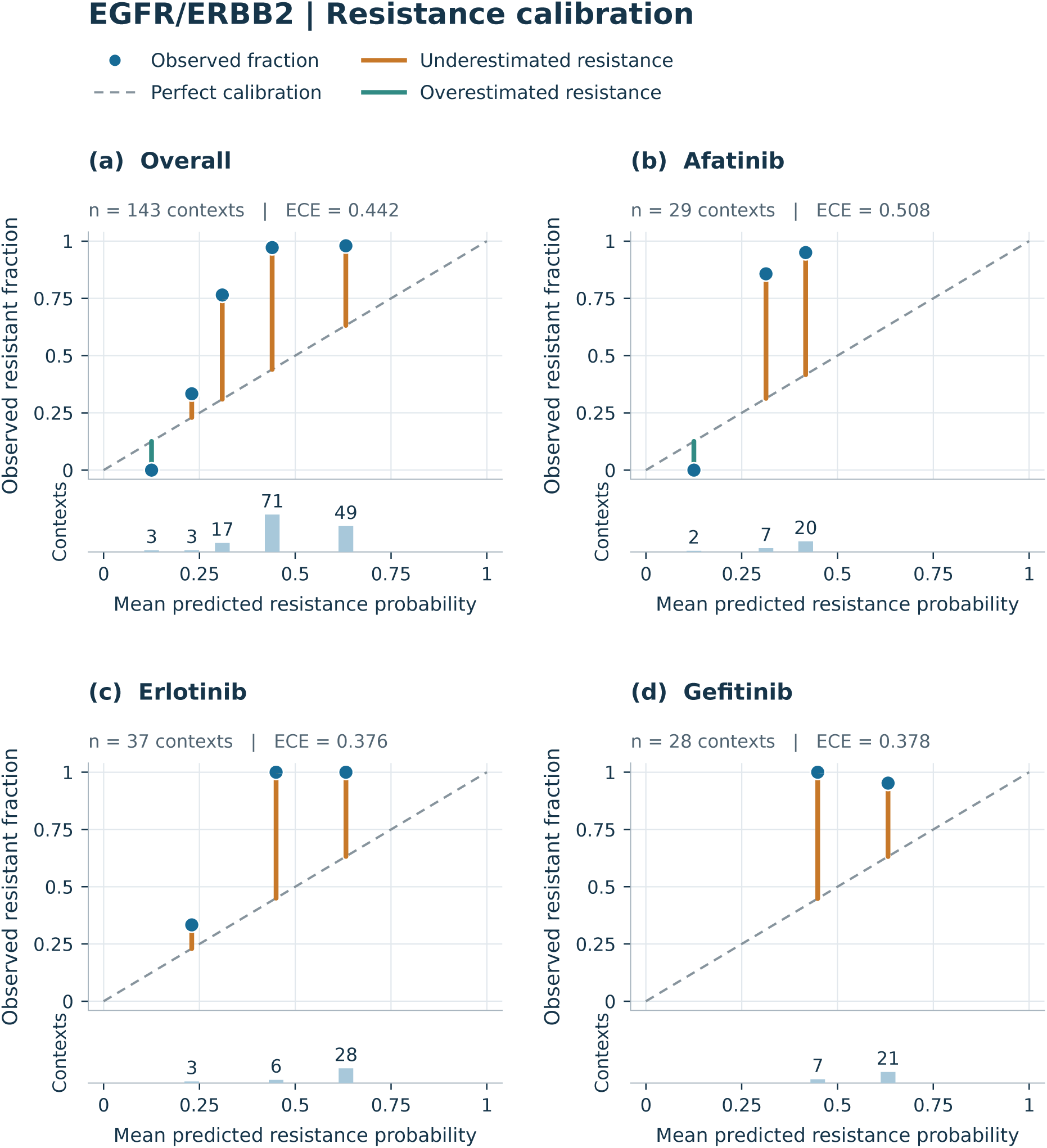
EGFR/ERBB2 classification calibration for the overall test set (a), afatinib (b), erlotinib (c), and gefitinib (d). Filled points show the observed resistant fraction against the mean predicted resistance probability in each non-empty bin. The dashed diagonal indicates perfect calibration. Vertical segments show underestimation (orange) or overestimation (teal) of the resistant fraction. Lower bars show bin counts on a common count scale within the figure; numerical labels give the exact counts. *n* denotes context rows in the panel, and ECE denotes expected calibration error. Empty bins are omitted.

**Figure S2:**
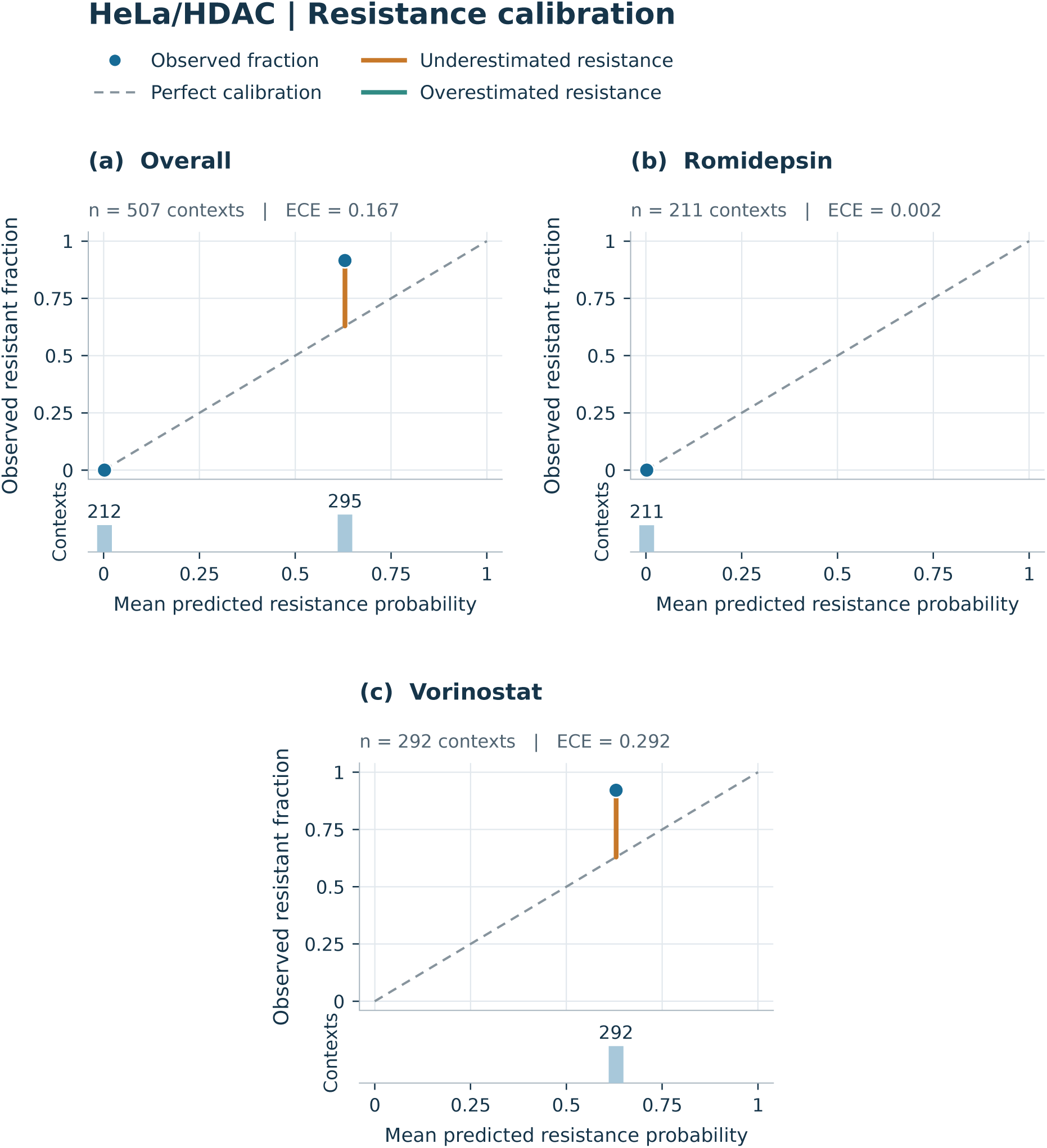
HeLa/HDAC classification calibration for the overall test set (a), romidepsin (b), and vorinostat (c). Filled points show the observed resistant fraction against the mean predicted resistance probability in each non-empty bin. The dashed diagonal indicates perfect calibration. Vertical segments show underestimation (orange) or overestimation (teal) of the resistant fraction. Lower bars show bin counts on a common count scale within the figure; numerical labels give the exact counts. *n* denotes context rows in the panel, and ECE denotes expected calibration error. Empty bins are omitted.

**Figure S3:**
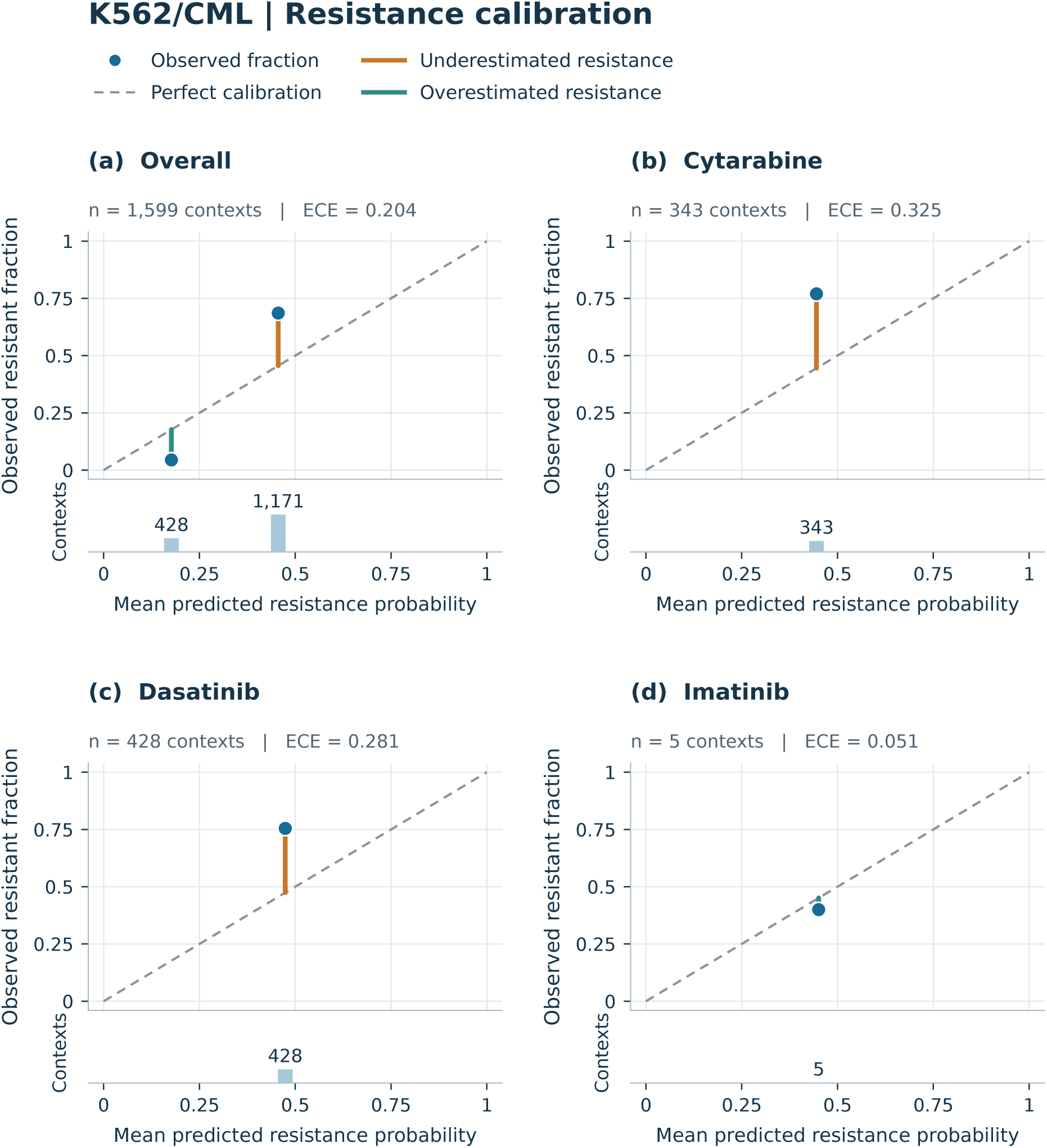
K562/CML classification calibration for the overall test set (a), cytarabine (b), dasatinib (c), and imatinib (d). Filled points show the observed resistant fraction against the mean predicted resistance probability in each non-empty bin. The dashed diagonal indicates perfect calibration. Vertical segments show underestimation (orange) or overestimation (teal) of the resistant fraction. Lower bars show bin counts on a common count scale within the figure; numerical labels give the exact counts. *n* denotes context rows in the panel, and ECE denotes expected calibration error. Empty bins are omitted.

## Notes

https://github.com/Moustafazn/PTM-BDL

### Competing Interest Statement

The authors have declared no competing interest.

